# Targeting the oncogene LSF with either the small molecule inhibitor FQI1 or siRNA causes mitotic delays with unaligned chromosomes, resulting in cell death or senescence

**DOI:** 10.1101/665570

**Authors:** Jennifer L.S. Willoughby, Kelly George, Mark P. Roberto, Hang Gyeong Chin, Patrick Stoiber, Hyunjin Shin, Chandra Sekhar Pedamallu, Scott E. Schaus, Kevin Fitzgerald, Jagesh Shah, Ulla Hansen

## Abstract

**Background:** The oncogene LSF (encoded by *TFCP2*) has been proposed as a novel therapeutic target for multiple cancers. LSF overexpression in patient tumors correlates with poor prognosis in particular for both hepatocellular carcinoma and colorectal cancer. The limited treatment outcomes for these diseases underscore the need for molecularly targeting novel mechanisms. LSF small molecule inhibitors, Factor Quinolinone Inhibitors (FQIs), have exhibited robust anti-tumor activity in multiple mouse models, with no observable toxicity.

**Methods:** Cell proliferation and cell cycle progression were analyzed after loss of LSF activity, using HeLa cells as a model cancer cell line responsive to FQI1. In addition, results were compared after treatment with either FQI1 or siRNA targeting LSF to test for biological specificity of targeting LSF by FQI1.

**Results:** Cellular phenotypes observed upon FQI1 treatment were due specifically to the loss of LSF activity, as siRNA targeting LSF produced highly similar phenotypes. Inhibition of LSF activity by either mechanism induced a strong delay prior to metaphase during progression through mitosis, with condensed, but unaligned, chromosomes. This mitotic disruption resulted in improper cellular division leading to multiple outcomes: multi-nucleation, apoptosis, and cellular senescence.

**Conclusions:** Specific inhibition of LSF by small molecules or siRNA results in mitotic defects, leading to cell death or senescence - consequences that are desirable in combating cancer. Taken together, these findings not only confirm that LSF is a promising target for cancer treatment, but also that FQIs are promising compounds for obtaining therapeutic effects for multiple LSF-driven cancers with unmet medical need.

## Background

LSF (encoded by *TFCP2*) is an evolutionarily conserved transcription factor that is normally expressed ubiquitously at low levels, but is significantly overexpressed in multiple specific cancers (1). This was initially shown in hepatocellular carcinoma cell lines and patient samples, in which levels of LSF in patient samples from multiple populations rise with increased stage and severity of disease (2–5). Furthermore, LSF is oncogenic for hepatocellular carcinoma, as it is sufficient to promote hepatocellular carcinoma tumor growth in mouse xenograft models (2, 3). In both colorectal cancer and hepatocellular carcinoma, patients with elevated LSF levels have significantly worse prognosis, with shorter median disease-free survival times than those with low LSF levels (4, 6). Finally, recent reports demonstrated that LSF can function as a co-activator for key transcription factors downstream of the Hippo and Wnt signaling pathways - YAP (5) and β-catenin (7) – both of which are widely accepted to contribute to liver proliferation and oncogenesis, as well as other cancer types.

Primary liver cancer and colorectal cancer are among the most common cancers worldwide (sixth and third, respectively), and represent leading causes of cancer mortality (second and fourth, respectively) (8–10). Hepatocellular carcinoma represents approximately 70-80% of primary liver cancer cases (9, 11). Although treatment options have improved, survival rates have only moderately increased. The two initial first-line FDA-approved therapies for late-stage hepatocellular carcinoma, Sorafenib and Regorafenib (multi-kinase inhibitors), demonstrate only modest improvement in patient survival rates (12, 13), and result in significant side effects and rapid development of drug resistance. A recent additional first-line treatment for unresectable HCC approved by the FDA, lenvatinib, demonstrates improvements in progression-free survival and objective response rate, although still limited improvement in survival (14). Thus, a large unmet medical need remains for hepatocellular carcinoma, as well as colorectal patient populations. Therapies directed to distinct molecular targets, ideally to which the cancer is oncogene addicted, have been promoted for mitigating these cancers (11).

A family of small molecule inhibitors of LSF, Factor Quinolinone Inhibitors (FQIs), was identified that inhibits the DNA binding and transcription activity of LSF, but not that of transcription factors from multiple other structural classes (15). Phenotypically, depletion of LSF by siRNA or FQIs inhibit growth of hepatocellular carcinoma or pancreatic cells *in vitro* (2, 5, 7, 15). They also inhibit hepatocellular carcinoma tumor growth *in vivo* in multiple mouse models, including a mouse endogenous liver tumor model (16). In all cases, inhibition of tumor growth occurred in the absence of toxicity, as assessed by liver injury markers, histopathology of tissues with rapid cell turnover, or blood cell counts (17). These results suggested that hepatocellular carcinoma cells are oncogene addicted to LSF (15, 18).

Oncogenic transcription factors are promising therapeutic targets given that they regulate tumorigenic pathways. However, transcription factors, in general, have been notoriously difficult to target with small molecule inhibitors as their DNA binding domains are commonly small and the proteins themselves are intrinsically disordered promiscuity (19). Identification of the transcription factor LSF as an oncogene and the significant inhibition of tumor growth upon LSF inhibition with no observed toxicity indicate that LSF holds considerable promise as a cancer therapeutic target (2, 15, 20). Targeting a transcription factor has been challenging, therefore validation of the biological specificity of the LSF inhibitors is essential. Here we demonstrate that the molecular and phenotypic consequences of knockdown of LSF with a specific siRNA are the same as treatment of cells with FQI1, therefore confirming that FQIs are highly specific in targeting this transcription factor.

The molecular mechanisms by which LSF promotes cancer cell survival have not been characterized in detail, although initial data indicated that FQIs induce a mitotic arrest in hepatocellular carcinoma cells (16). Clarifying the pathways by which inhibition of LSF leads to cell death is important to support the candidacy of FQIs as a molecular therapy. Cell cycle analysis by flow cytometry and time-lapse microscopy revealed mitotic defects including mitotic delays with condensed, but unaligned chromosomes, leading to increased time in mitosis, defective cell division, multi-nucleation, and apoptosis. In addition, loss of LSF activity induced senescence in a sub-population of cells in a dose-dependent manner. Senescence, as well as mitotic arrest and apoptosis, are all desirable outcomes for a cancer chemotherapeutic.

## Methods

### Preparation of FQI1

FQI1 was synthesized as previously described (15). FQI1 was dissolved in analytical grade DMSO (Sigma). The final DMSO concentration added to the cells was 0.5%.

### Cell lines and synchronization

HeLa cells (gift from Devanand Sarkar, Virginia Commonwealth University) were cultured at 37°C in 10 % CO_2_ in DMEM (Corning Cellgro) supplemented with 10% Fetal Bovine Serum (FBS; Atlanta Biologicals). Validation of HeLa cells was performed both by PCR of genomic DNA to confirm presence the E7 region of HPV18 and RNA-seq analysis confirming expression of the precise regions of the HPV18 genome that are present in HeLa cells. Prior to harvesting, cells underwent 4-7 passages. Cells tested negative for mycoplasma genus by PCR (Charles River Research). For synchronization using a double thymidine block protocol, cells were treated with 2 mM thymidine (Sigma) in complete medium for 18 hours, and then released into complete medium for 6 hours followed by incubation in 2 mM thymidine in complete medium for a second 18-hour incubation. For release from the G1/S block, cells were transferred into complete medium. The single thymidine block involved a single 24 hour incubation in 2 mM thymidine and release into complete media. As indicated, the release medium also contained 20 μM of thymidine.

### siRNA transfection

siRNAs were designed and synthesized at Alnylam Pharmaceuticals, Inc. siRNA sequences used: LSF Sense: 5’ GUGUGAUGUUUAACAGGAATT 3’; LSF Antisense: 5’ UUCCUGUUAAACAUCACACTT 3’; LBP1A Sense 5’ UUUCAGGUGCCGACUUAUUTT 3’; LBP1A Antisense: 5’ AAUAAGUCGGCACCUGAAATT 3’. The siRNAs were stabilized using certain chemical modifications as previously described allowing durable knockdown (21, 22). The siRNA control was a sequence targeting RNA encoding firefly luciferase and was, therefore, non-targeting in the cells utilized for these studies. Cells were transfected using RNAimax (Life Technologies) according to manufacturer’s instructions. Transfection efficiency was measured by fluorescent microscopy 24 hours post transfection by cellular uptake of the Cy3 labeled control siRNA, and was determined to be >90%. For all siRNA experiments, the initial thymidine block was started 24 hours after transfection of the siRNA.

### Time-lapse microscopy

In order to image cell cycle progression for HeLa cells, retroviral Packaging Cells (GP2-293; Clontech) were transfected with pVSV-G (Clontech) and a pBABE vector containing both a gene for YFP-tagged histone H2B protein and for a gene encoding G418 resistance. The virus-containing supernatant was collected for transduction of HeLa cells and the population of resistant cells was selected with G418 (Gibco).

For time-lapse microscopy of siRNA-treated cells, the HeLa cells expressing H2B-YFP were transfected and synchronized with a single thymidine block and release. After release, cells were imaged in CO_2_ independent medium (Leibovitz’s L-15 without phenol red) on a Nikon TA10 Eclipse with a 20X objective at 37°C. For time-lapse microscopy of FQI1-treated cells, HeLa cells expressing H2B-YFP were treated with either vehicle or 0.9, 1.8, or 3.6 µM FQI1 in CO_2_ independent medium (Leibovitz’s L-15 without phenol red). Cells were imaged immediately on a Nikon TA10 Eclipse with a 20X objective at 37°C. Images were acquired every four minutes at 7-10 positions per sample, over a five-to eleven-hour time span. Length of mitosis was measured from nuclear envelope breakdown to anaphase. Nuclear envelope breakdown was identified as the first image displaying disordered, condensed chromosomes. Anaphase was identified as the first image showing sister chromatid separation (for normal anaphases) or showing a furrow beginning to form over the chromosomes. For experiments, 100 to 101 cells were examined per condition.

### Cell flow cytometry

Cells were harvested, washed, and fixed with ethanol. Cells were stained with the Guava cell cycle reagent (EMD Millipore) according to manufacturer’s instructions. Fluorescence was analyzed on a BD Dickenson FACS Calibur.

### Immunoblotting

Cells were lysed in RIPA buffer (125 mM Tris HCl, 150 mM NaCl, 0.1% NP-40, 1.0% Sodium deoxycholate, 1.0% SDS, pH 7.6) containing ROCHE protease cocktail phosphatase inhibitors (Sigma Aldrich 4693159001) at the manufacturer’s recommended concentrations. Lysates were electrophoresed through 4-20% Mini-PROTEAN^®^ TGX™ Precast gradient gels (Bio-rad). The proteins were transferred to a PVDF membrane, and membranes were incubated for 1 hour in Odyssey Blocking buffer (LI-COR Biosciences cat# 927-40000). Primary antibodies included Aurora Kinase B (Abcam AB2254), Cdc20 (Abcam AB26483), Cyclin B1 (AB72), LBP-1a (ABE181), LSF (Abcam ABE180), phosphorylated Histone 3 Serine 10 (Abcam ab5176), phosphorylated Histone 3 Serine 28 (Abcam, ab5169), and α-Tubulin (Sigma, 10002). Secondary antibodies were from LI-Cor, Inc. and included donkey anti-mouse IR800 (926–32212), donkey anti-rabbit IR800 (926–32213), goat anti-rabbit IR680 (926–68073), and goat anti-mouse IR680 (926–32214). PVDF membranes were imaged using the LI-COR Odyssey (23). Infrared detection quantitated each band on an individual pixel basis using western analysis tools in the Image Studio program.

### Gene expression determination

For most experiments, RNA was isolated using the Qiagen RNAeasy kit following the manufacturer’s instructions. cDNA was generated using a Reverse Transcription kit from Applied Biosystems (4368814). Probes for RNA quantification were acquired from Life Technologies with the Taqman gene expression system (Life Technologies). Target gene expression was normalized to a ubiquitous control (*GAPDH*) utilizing a dual label system. Cp values were measured using a Light Cycler 480 (Roche). The following probes were used: (*AURKB*) HS009645858 M1, (*CDC20*) HS00426680 M1, (*UBP1*, which encodes LBP1A) HS00232691 M1m, (*TFCP2*, which encodes LSF) HS00232185 M1, (*MAD2L1*) HS00365651 M1, and (*GAPDH*) 4333764F.

### Cellular senescence measurement

HeLa cells were synchronized with a double thymidine block with either FQI1 treatment or LSF knockdown. The cells were stained for β-galactosidase using the activity kit (Kit 9860S) from Cell Signaling Technologies according to the manufacturer’s protocol. Following the overnight incubation, cells were imaged on a phase Axiovert 40 CFL (Zeiss) microscope. The number of blue-staining cells was quantified, irrespective of the intensity of the signal, in comparison to the number of cells lacking blue staining.

### Statistical analysis

Statistical significance was determined using a two-tailed Student T Test (paired tests unless otherwise stated); *P<0.05, **P<0.01, ***P<0.001, ****P<0.0001 using Prism GraphPad. Pearson correlation coefficient was determined using Excel.

## Results

### Chemical inhibition of LSF induces a delay in mitotic progression with condensed but unaligned chromosomes

In asynchronous populations of hepatocellular carcinoma cells, the LSF small molecule inhibitors (FQIs) resulted in cells delayed with G2/M (“4n”) DNA content (16). In order to characterize this cell cycle-related phenotype in detail, we utilized HeLa cells, which are similarly sensitive to FQI1 treatment and readily able to be synchronized for in depth cell cycle analysis. First, the dose- and time-dependence of this phenotype was examined, using cellular DNA content as a readout. Cells were synchronized at G1/S with a double thymidine block (Fig. 1A) and analyzed for cellular DNA content throughout the subsequent cell cycle. At 1.8 µM, FQI1-treated cells were initially delayed in returning from G2/M to G1, remaining with 4n DNA content, compared to control cells that had re-entered G1 (Fig. 1B, 8.5 h), an observation consistent with previous studies (16). Following this delay, a mix of phenotypic outcomes was observed at 20 hours after release from G1/S, with FQI1-treated cells having divided to a 2n DNA content, initiated cell death pathways (subG1 DNA content), or retained their duplicated DNA content. At the highest FQI1 concentration tested (3.6 µM), cells were also initially delayed with 4n DNA content, but a large fraction of the population converted to subG1 DNA content by 20 hours post release from the G1/S block (Fig. 1B).

**Figure 1.**
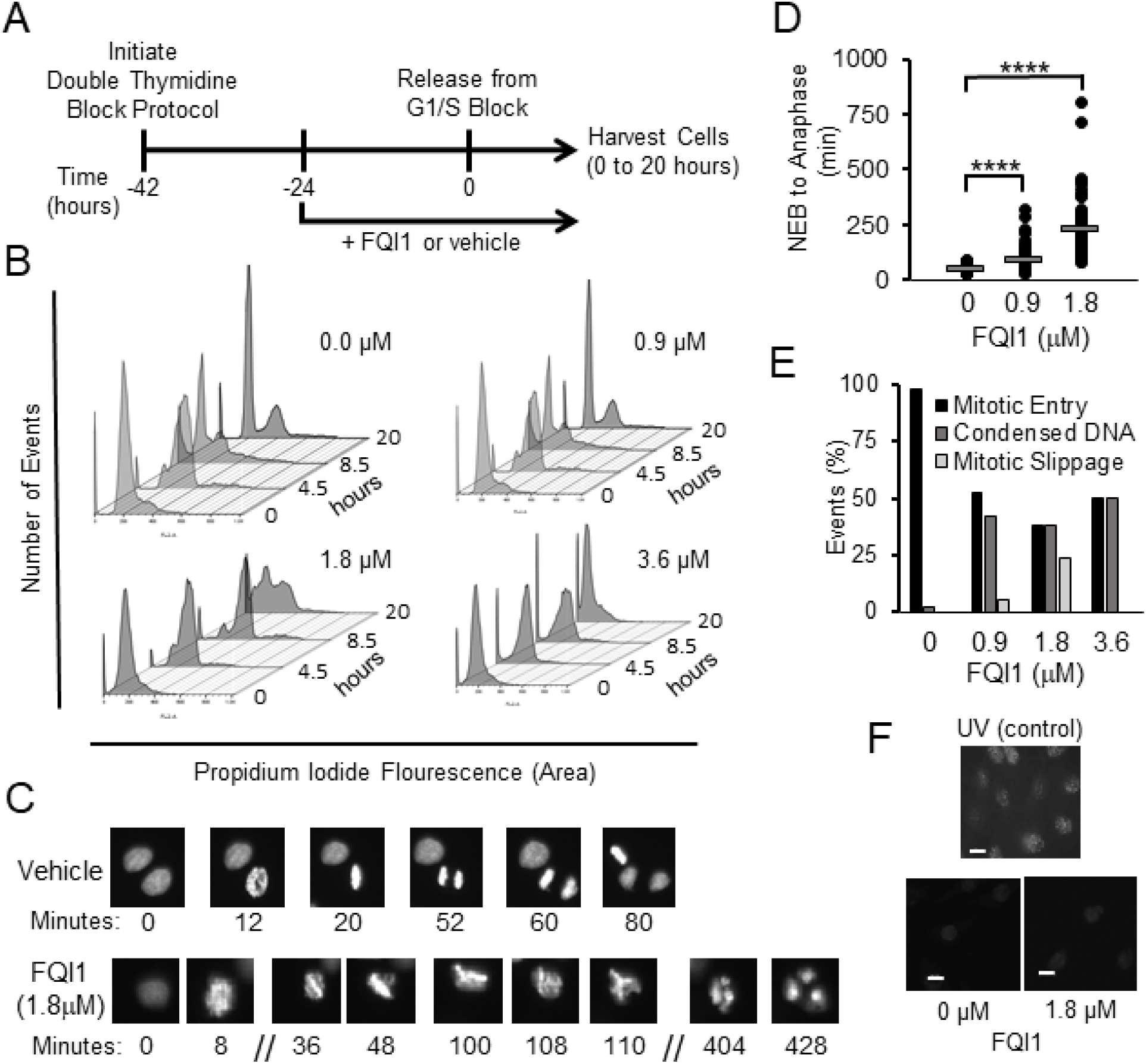
FQI1-treated HeLa cells exhibit mitotic defects. **A.** Schematic of experimental protocol. FQI1 or vehicle was added to HeLa cells during synchronization to the G1/S border using a double thymidine block. Cells released from the block in the presence of the FQI1 or vehicle, plus 20 µM of thymidine, were harvested at multiple times during progression through the cell cycle. **B.** At the indicated time points following release from the G1/S block with 0, 0.9, 1.8, or 3.6 µM of FQI1, cells were analyzed for DNA profiling by flow cytometry. Data are representative of at least three independent experiments. **C.** Representative time-lapse images of individual cells treated with vehicle or 1.8 µM FQI1. Numbers represent the time (in minutes) for one particular cell in the image from nuclear envelope breakdown (designated as time=0 for that cell). **D.** Quantitation of mitotic time from nuclear envelop breakdown (NEB) to anaphase for the population of asynchronous cells during treatment for approximately 16 hours with FQI1 or vehicle. Mitotic times (mean time in minutes +/− standard error of the mean, n) for vehicle, and 0.9 or 1.8 µM FQI1 treatments were: 48.7 +/− 1.5, 104; 84.5 +/− 4.9, 104; and 228 +/− 15, 77; respectively. Mitotic time for cells treated with 3.6 µM was not quantifiable, as those cells that entered mitosis during the imaging period never reached anaphase or nuclear division. **E.** Quantitation of cellular events at increasing concentrations of FQI1 during the time lapse microscopy, including percentage of cells that entered mitosis but were delayed with condensed, but unaligned chromosomes, and the percentage that apparently underwent mitotic slippage with formation of multiple (>2) nuclei. 120-140 cells were analyzed for each concentration of FQI1, including vehicle alone. **F.** Bottom: γ-H2AX staining of HeLa cells treated with vehicle or 1.8 µM FQI1. Top: Representative image of UV-treated HeLa cells as a positive control. All images were taken at the same intensity and are representative of two independent experiments. Scale bars: 20 µm.

Given limitations in interpretation of population-wide data, it was critical to analyze the phenotype(s) on a per cell basis upon inhibition of LSF. Using HeLa cells stably expressing fluorescently labeled histone H2B, chromosomal DNA was visualized as cells passed through mitosis by time-lapse microscopy. Asynchronous H2B-YFP-expressing cells were treated with increasing concentrations of FQI1. At 1.8 µM (Fig. 1C) mitotic progression was delayed with condensed, unaligned chromosomes. Cells subsequently appeared to exit mitosis without proper chromosome segregation, resulting in a multinuclear (4n) G1 state (Fig. 1C) (16). In contrast, cells treated with vehicle progressed through mitosis normally (Fig. 1C and D). Mitotic time (time from nuclear envelope breakdown (NEB) to anaphase) was dose-dependent, increasing with increasing concentrations of FQI1 (Fig. 1D). At the lower concentrations of FQI1, cells exited mitosis aberrantly after the mitotic delay (Fig. 1C, D and E). However for cells treated with 3.6 µM FQI1, mitotic time was indeterminable (Fig. 1E); cells arrested with condensed but unaligned chromosomes but never exited mitosis, either normally or aberrantly, throughout the 640 minutes of imaging (Supplementary Fig. S1A).

In some cell lines, LSF is necessary for upregulation of thymidylate synthase expression and therefore efficient transition through S phase (24). Since S phase defects can ultimately lead to mitotic defects, DNA damage was monitored by measuring phosphorylated H2AX (γ-H2AX) chromosomal foci. Cells treated with 1.8 µM FQI1 or vehicle during synchronization were analyzed for γ-H2AX by immunofluorescence 8 hours after release from the G1/S block (Fig. 1F). UV-irradiated cells, a positive control, demonstrated extensive γ-H2AX staining. In contrast, only comparable, low levels of phosphorylated H2AX were visualized in both the vehicle- and FQI1-treated cells, consistent with low levels of DNA damage known to occur in cancer cells (25). This was consistent with our previous studies suggesting that adequate thymidylate synthase expression in tumor cells was likely achieved even with reduction in LSF activity (15, 26).

Overall, upon LSF activity inhibition, we observed defects in chromosome alignment and segregation, resulting in mitotic delay and multi-nucleation. The lack of elevated DNA damage signals supports the hypothesis that FQI1-mediated mitotic defects are due to a direct requirement for LSF in regulating proper progression through mitosis, and in particular in progressing to metaphase, in which the condensed chromosomes are fully aligned.

### LSF small molecule inhibition during cell synchronization reduced expression of mitotic regulators

The lack of chromosomal alignment is not a phenotype expected to be caused by upregulation of cyclin B expression, which was previously suggested from experiments in asynchronous cells to be a major cause of the FQI1-mediated mitotic delay (16). Thus, we investigated the effects of FQI1 treatment on the expression of a number of mitotic regulators, using synchronized cells to more carefully assay cell cycle expression. HeLa cells were synchronized with a double thymidine block (Fig. 2A). RNA levels were measured in vehicle-treated cells at the G1/S border (0 hours) and as cells progressed to or just through mitosis (8 hours), as demonstrated by DNA profiling (Fig. 1B) and fluorescence microscopy (Supplementary Fig. S1B). Results are shown for the mitotic regulators Aurora kinase B (*AURKB*) and Cyclin Division Cycle 20 (*CDC20*) (27, 28), as expression levels of these genes were altered by FQI1 (see below). In the untreated, control cells, mitotic expression of *CDC20* was elevated approximately 6-fold compared to RNA levels at G1/S (Supplementary Fig. S1C), as expected. However, *AURKB* RNA levels in vehicle-treated cells increased only 1.2 fold in mitosis, consistent with dysregulated expression in these cancer cells. Incubation with 1.8 µM FQI1 during the synchronization protocol resulted in reduction of both *AURKB* and *CDC20* RNA levels compared to the control cells 8 hours post release (Fig. 2B). Consistent with transcript reduction, AURKB and CDC20 protein levels were also reduced in a dose-dependent manner (Fig. 2C and 2D), whereas LSF protein levels were unchanged, as expected (Fig. 2C). The impact of the downregulation of AURKB was tested by monitoring phosphorylation of an AURKB substrate. Phosphorylation of Histone 3 on Serine 10 (29) was reduced by FQI1 in a dose-dependent manner (Fig. 2C and 2D).

**Figure 2.**
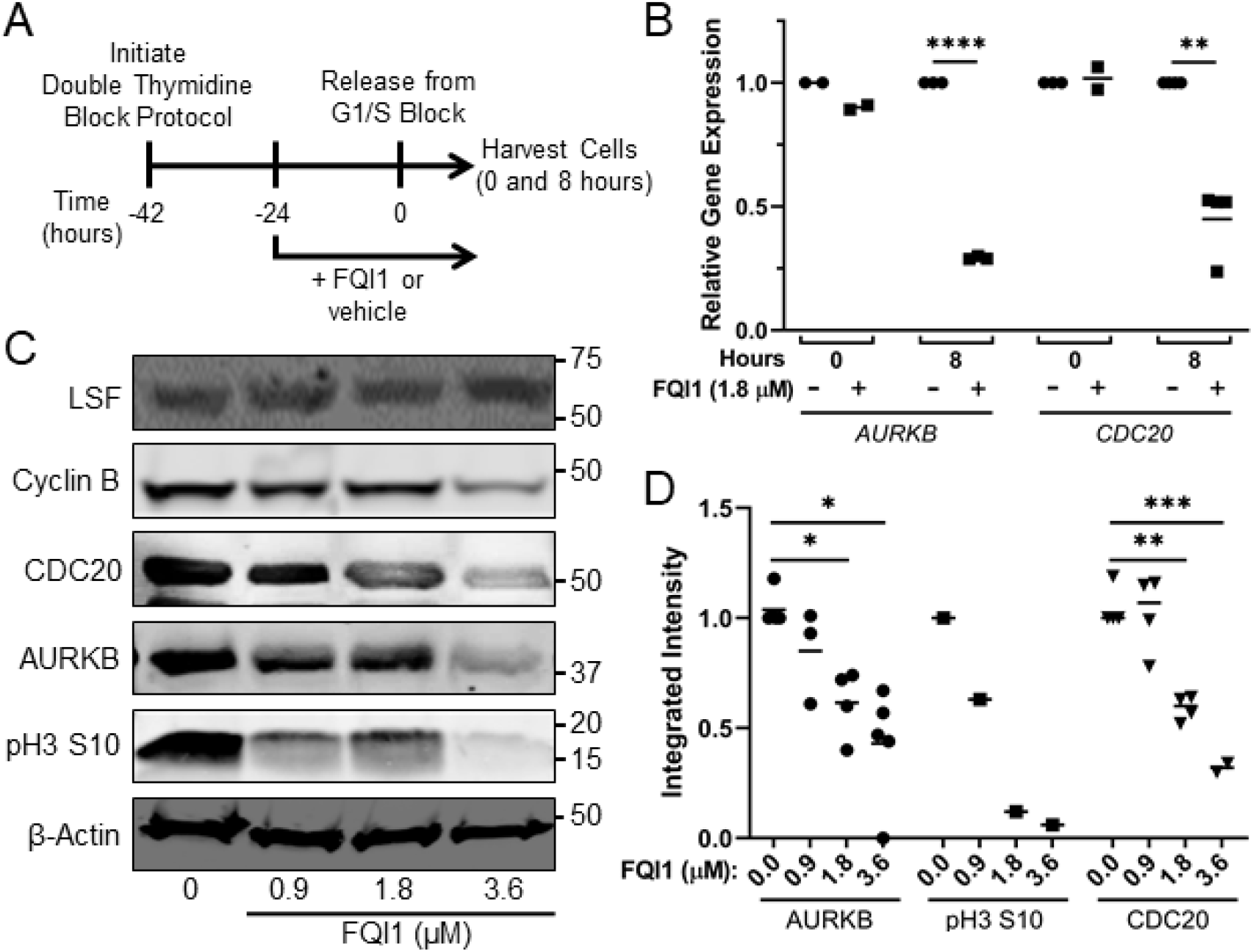
FQI1 treatment diminished expression of mitotic regulators. **A.** Schematic of experimental protocol. FQI1 or vehicle was added to HeLa cells during synchronization to the G1/S border using a double thymidine block. Cells were released from the block, including addition of 20 µM of thymidine, for subsequent analyses. **B.** Lysates from cells treated with vehicle or 1.8 µM FQI1 were harvested at release from the G1/S block (0 hours) or 8 hours post release and analyzed for *AURKB* or *CDC20* RNA levels, as normalized to levels of *GAPDH* RNA. Data points and means are plotted relative to the expression from vehicle treated cells at each time point and are derived from 2-4 independent experiments. ** p=0.0045; **** p<0.0001. **C.** Representative immunoblots for the indicated proteins in lysates harvested 8 hours post release from a G1/S block, after treatment with increasing concentrations of FQI1. Molecular weight markers are indicated on the right side. **D.** Quantitation of the protein levels from immunoblots (e.g. panel C). Protein levels were normalized to those of β-actin. Data points and means are from 2-5 independent experiments. *p = 0.037 (FQI1=1.8 µM), 0.015 (FQI1=3.6 µM); **p=0.0048; ***p=0.0005 (unpaired T test).

In stark contrast to the previous results in asynchronous cells (16), in the synchronized cells treated with 1.8 µM FQI1, Cyclin B protein levels were similar to those in control cells, and certainly not increased (Fig. 2C), despite obvious delays in mitotic progression at this concentration (Figs. 1B-C). Furthermore, at the higher concentrations of FQI1 during the synchronization procedure, Cyclin B levels were actually downregulated (Fig. 2C). Thus the cause of the mitotic defects cannot be elevated cyclin B levels.

Because cyclin B is required for mitotic entry, however, the lower levels of cyclin B suggested that cells treated with higher concentrations of FQI1 during synchronization were not efficiently proceeding into the final mitosis after release from the final thymidine block (see also below). This is consistent with the bulk cellular DNA profiling curves (Fig. 1B), since this analysis cannot distinguish between “2n” as early G1 *versus* G1/S or “4n” as G2 *versus* M. This complicates straightforward interpretation of the bulk expression results, with ambiguity as to whether decreased levels of AURKB and/or CDC20 were the cause of the mitotic cell cycle defects, or simply the consequence of cell cycle defects resulting from lower LSF activity during the first passage through mitosis in the synchronization procedure.

For a stringent examination of whether diminished *AURKB* and *CDC20* gene expression resulted from lack of cell cycle progression of LSF inhibited cells or from diminished expression of these genes in mitosis in the presence of FQI1, we analyzed RNA in synchronized, LSF-inhibited cells only from cells demonstrably in mitosis, isolated by standard mitotic shakeoff methodology. A reproducible decrease in *CDC20* (Supplementary Fig. S2B), but not *AURKB* (Supplementary Fig. S2A), RNA was observed in this experiment. We also sought to identify candidate LSF target genes by identifying binding sites for LSF near the genes. Given the lack of a sufficiently robust antibody against LSF for chromatin immunoprecipitation (ChIP), a stable HEK cell line inducibly expressing HA-tagged LSF (15) was used for the ChIP-sequencing analysis. Multiple HA-LSF binding peaks were observed around the *AURKB* gene (Supplementary Fig. S2C), and binding of LSF was validated both at the *AURKB* promoter and around 3000 bp upstream of the transcription start site by quantitative PCR (Supplementary Fig. S2D). In contrast, no HA-LSF binding peaks were observed within 20 kb of the *CDC20* gene. Taken in combination, whether LSF activates *AURKB* expression in these, or other, cells remains unresolved. The mitotic shakeoff experiment suggests that LSF may regulate *CDC20* expression, either from distant binding sites, or indirectly. Global gene expression data from cells treated with FQI1 only between G1/S and mitosis did not identify dysregulation of RNA encoding any other mitotic regulators (30).

Despite not pinpointing mitotic genes directly transcriptionally regulated by LSF, these results did provide molecular biomarkers in this synchronized cell system for responsiveness to the LSF inhibitor FQI1.

### RNAi mediated knockdown of LSF phenocopies inhibition of LSF with the small molecule inhibitor FQI1

Specificity of small molecule inhibitors to their intended target is a key requirement so that biological consequences of inhibitor effects can be mechanistically attributed to the target of interest. Knowledge of specificity is of even more importance in developing such inhibitors for use in the clinics. FQI1 inhibits LSF DNA-binding and protein-binding activities, whereas it does not impact activity of a number of other transcription factors, both with disparate and similar structural domains (15, 31). However, in order to demonstrate that the overall cellular consequences of FQI1 treatment were specific consequences due to inhibiting LSF, a direct comparison with specific removal of LSF was required. Although LSF has a long half-life, of approximately 24 hours (32), we identified an siRNA that resulted in robust and durable knockdown of LSF (Fig. 3B and C, Supplementary Fig. S3A-C). In addition, since certain siRNAs can cause nonspecific reduction in mRNA encoding MAD2 (33), which controls the spindle assembly checkpoint in mitosis, we verified that the selected siRNA targeting LSF did not inadvertently reduce *MAD2L1* transcript levels (Supplementary Fig. S3D).

**Figure 3.**
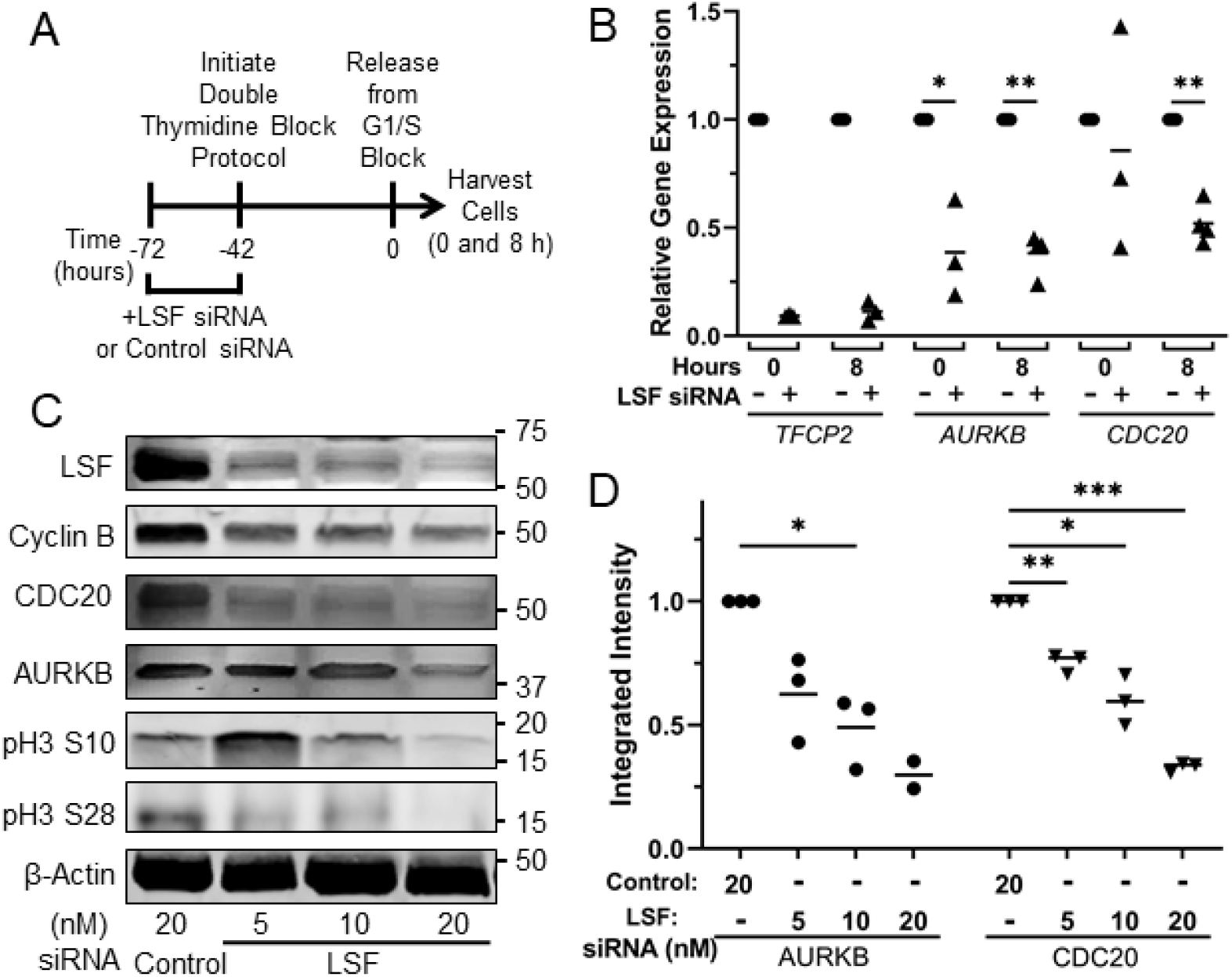
RNAi mediated knockdown of LSF reduced expression of mitotic regulators. **A.** Schematic of experimental protocol. siRNAs targeting LSF or a non-expressed target were transfected into HeLa cells at the indicated concentrations, followed by synchronization of cells and release from the G1/S block. **B.** Cells synchronized during treatment with 20 nM of LSF (+) or control siRNA (-) were harvested for RNA at 0 or 8 hours after release from the final G1/S block. *TFCP2* (which encodes LSF), *AURKB*, and *CDC20* RNA levels were measured and normalized to those of *GAPDH* from the same time point. The relative gene expression levels are reported as the fraction of the RNA levels in the control siRNA-treated cells at 0 hours. Data points and means are from 3-4 independent experiments. *p=0.042; **p=0.0010 (*AURKB*), p=0.0019 (*CDC20*). **C.** Representative immunoblots of the indicated proteins or protein modifications are shown for lysates harvested at 8 hours post release from the final G1/S block. Molecular weight markers are indicated on the right side. **D.** Quantification of immunoblots of AURKB and CDC20 (e.g. panel C). Each target protein was normalized to the level of β-actin in the same sample. Data points and means are from 2-3 independent experiments. *p=0.027 (AURKB), p=0.021 (CDC20); **p=0.0087; ***p=0.0003.

In order to compare downstream biomarkers from FQI1 and LSF siRNA treatments, *AURKB* or *CDC20* RNA levels were measured following RNAi mediated knockdown of either LSF or a non-expressed control. Cells were transfected with siRNAs, to initiate protein knockdown, 30 hours prior to synchronization. RNA and protein expression were analyzed at two time points - when control cells were arrested at G1/S (0 hours) following the synchronization protocol and when these cells were largely in mitosis after release from the block (8 hours) (Fig. 3A, Supplementary Fig. S4A). For ease of comparison, RNA levels were plotted relative to the level in the control siRNA sample at each time point. At 20 nM siRNA, significant knockdown of both LSF-encoding RNA (*TFCP2*, Fig. 3B) and protein (Fig. 3C) were achieved over this time course. Consistent with the results generated with the LSF small molecule inhibitor at 8 hours after G1/S release (Fig. 2B), *AURKB* and *CDC20* RNA levels were significantly reduced upon knockdown of LSF (Fig. 3B). Immunoblotting of lysates harvested at the approximate time of mitotic entry of the control cells (8 hours) confirmed a dose-dependent reduction in AURKB and CDC20 protein levels after siRNA-mediated knockdown of LSF (Fig. 3C and D), consistent with the findings upon inhibition of LSF with FQI1 (Fig. 2C and D). As expected, phosphorylation of AURKB substrates Serine 10 and 28 of histone H3 (29, 34) was reduced (Fig. 3C). Finally, as with FQI1 treatments, Cyclin B levels were also reduced in a dose-dependent manner. Thus, LSF siRNA phenocopied the molecular consequences of FQI1 on protein expression, whether due to direct transcriptional effects, and/or consequences of cell cycle dysregulation.

To determine whether LSF knockdown resulted in similar mitotic phenotypes to those observed with FQI1, synchronized YFP-H2B-expressing HeLa cells were transfected with siRNAs targeting LSF or a non-expressed control. A single thymidine block protocol was sufficient for synchronization (Fig. 4A), as mitotic progression is viewed on a cell-by-cell basis. Representative time-lapse images of cells treated with the highest concentration (20 nM) of either LSF targeting siRNA or control siRNA highlight dramatic changes in mitotic progression. Control cells exhibited progression through normal mitotic phases in a timely manner (Fig. 4B). However, cells with diminished LSF levels exhibited an extensive delay with condensed chromosomes that never achieved stable alignment, generally followed by defective cellular division and multinucleation (Fig. 4B). In addition, some cells remained in mitosis with condensed chromosomes throughout the entire time lapse analysis. Quantitation documented that mitotic time was dramatically increased when LSF levels were reduced (Fig. 4C). We note that 5 and 10 nM of LSF siRNA resulted in slightly longer times for mitotic progression, which is likely due to the inability of a number of cells at the 20 nM LSF siRNA treatment (the most perturbed cells) to even enter mitosis. When siRNA-transfected cells were imaged by time lapse microscopy after a double thymidine block, there was an inverse correlation between higher levels of LSF knockdown and the number of cells capable of entering mitosis (Supplementary Fig. S4B).

**Figure 4.**
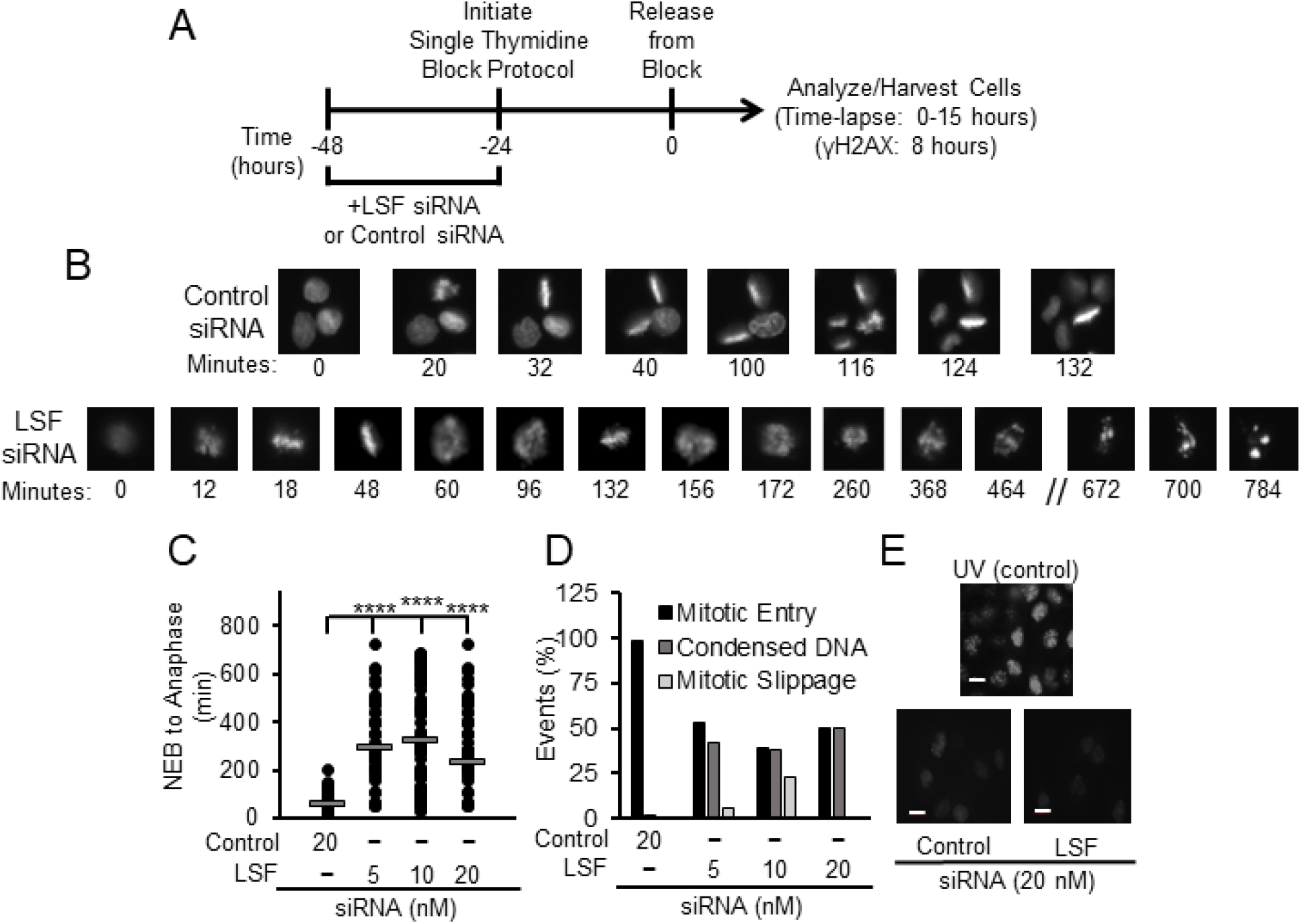
LSF knockdown in HeLa cells results in mitotic defects. **A.** Effects of LSF-specific siRNA on synchronized HeLa cells expressing YFP-labeled H2B were analyzed utilizing time-lapse microscopy. Schematic of experimental protocol for panels B-E. **B.** Representative images of cells treated with 20 nM of either control siRNA (top) or LSF siRNA (bottom). Numbers represent the time (in minutes) for one particular cell in the image from nuclear envelope breakdown (designated as time=0 for that cell). **C.** Quantitation of mitotic time from nuclear envelope breakdown (NEB) to anaphase for a population of cells treated with control siRNA or siRNA targeting LSF. Mitotic times (mean time in minutes +/− standard error, n) for 20 nM control siRNA, and 5, 10, or 20 nM LSF siRNA were: 57.9 +/− 2.8, 101; 296 +/− 16, 77; 324 +/− 25, 48; and 235 +/− 16, 84; respectively. Error bars indicated standard error of the mean, based on the number of cells analyzed in a single experiment. **D.** Quantitation of cellular events at increasing concentrations of LSF siRNA during the time lapse microscopy, including the percentage of cells that entered mitosis but were delayed with condensed but unaligned chromosomes, and the percentage that exited mitosis, but with multinucleation. The control had neither of these phenotypes among the cells counted (~100 per group). **E.** Bottom: γ-H2AX staining of HeLa cells treated with 20 nM control or LSF siRNA. Top: Representative image of UV-treated HeLa cells as a positive control. All images were captured at the same intensity. Scale bars: 20 µm.

By histone H2B fluorescence, the most striking mitotic outcome for individual cells treated with LSF siRNA after the extended delay in mitosis appeared to be mitotic slippage (Fig. 4B). This was confirmed by immunofluorescence of synchronized cell populations, co-stained for DNA and α-tubulin (Supplementary Fig. S5A-B). Quantitation of these immunofluorescence data demonstrated significant increases in cells with condensed, but nonaligned chromosomes, incomplete cytokinesis, and multinucleation upon LSF knockdown. These phenotypes mimicked those observed with FQI1 treatment (Supplementary Fig. S5C-D). Finally, both types of treatments yielded mitotic cells with cellular protrusions (Supplementary Fig. S5E).

Also consistent with the results from FQI1 treatment, knockdown of LSF did not induce phosphorylated H2AX (γ-H2AX) foci, as monitored the beginning of mitosis in a synchronized cell population (Fig. 4E). This result suggests that the effects of inhibiting LSF on mitotic progression are not due to defects induced indirectly in S phase.

LSF has a widely expressed paralog, LBP1A (35), whose activity is also inhibited by FQI1 (T. Grant, unpublished observations). Unlike LSF, however, LBP1A has not yet been implicated in cancer (1, 2), which is consistent with our results that knockdown of LSF alone caused the same molecular and phenotypic outcomes as those caused by FQI1. Nonetheless, to determine the potential contribution of LBP1A to FQI1 outcomes, we investigated the mitotic phenotypes upon LBP1A siRNA treatment. An siRNA was identified that resulted in robust and durable knockdown of LBP1A (Supplementary Fig. S6A-B), with no reduction in *MAD2L1* transcript levels (Supplementary Fig. S6C). Knockdown of LBP1A did reduce overall cell proliferation to some extent, but much less so than did knockdown of LSF (Supplementary Fig. S7A). Despite that proliferative effect, LBP1A knockdown did not significantly diminish cell viability, in stark contrast to consequences of LSF knockdown and FQI1 treatment (Supplementary Fig. S7B-C). Furthermore, LBP1A knockdown did not observably inhibit mitotic progression as measured by cellular DNA profiling (Supplementary Fig. S7D). However, time-lapse microscopy did detect a subtle (1.5–fold) increase in mitotic time upon LBP1A knockdown alone (Supplementary Fig. S7E), consistent with the diminished cell proliferation. No abnormal mitotic phenotypes were observed either by time-lapse or immunofluorescent microscopy. Given the minimal consequences upon inhibiting LBP1A, we conclude that inhibition of LSF activity is what drives the dramatic FQI1-mediated mitotic defects.

### Induction of cellular senescence following inhibition of LSF

In addition to mitotic delay resulting from LSF inhibition, some cells undergoing synchronization while inhibiting LSF were arrested at other points in the cell cycle, as suggested above by the reduction in cyclin B expression. Cellular DNA profiling of FQI1- or LSF siRNA-treated cells being synchronized with a double thymidine block, captured cells that no longer progressed from the 2n state into S phase upon release from the G1/S block (Fig. 1B, Supplementary Fig. S4A), and time-lapse microscopy showed that a considerable fraction of the cells treated with LSF siRNA during a double thymidine block never entered mitosis during 10-12 hours after release from the G1/S block (Supplementary Fig. S4B). Mitotic defects, caused by multiple distinct insults, can lead to senescence after G1 re-entry with either 2n or 4n DNA content (36, 37). Thus, we hypothesized that mitotic defects from decreasing LSF levels or activity during previous cell divisions resulted in senescence. To test this hypothesis, cells were synchronized as before by a double thymidine block in the presence of FQI1, or LSF siRNA, and analyzed for senescence by monitoring β-galactosidase activity at low pH (38) at a time point when control cells entered mitosis. Both reduction in LSF levels and inhibition of LSF activity resulted in significantly greater numbers of β-galactosidase-positive cells (stained blue) compared to the respective controls (Fig. 5A-B). Overall, there was a 3-to 5-fold increase in senescent cells with increasing amounts of LSF inhibition, although treatment with 0.9 μM FQI1 was not sufficient to induce senescence (Fig. 5C-D). These data show that inhibition of LSF can result in senescence of cancer cells, and support the hypothesis that reduced LSF levels or activity during previous cell cycle(s) can predispose cells to senescence.

**Figure 5.**
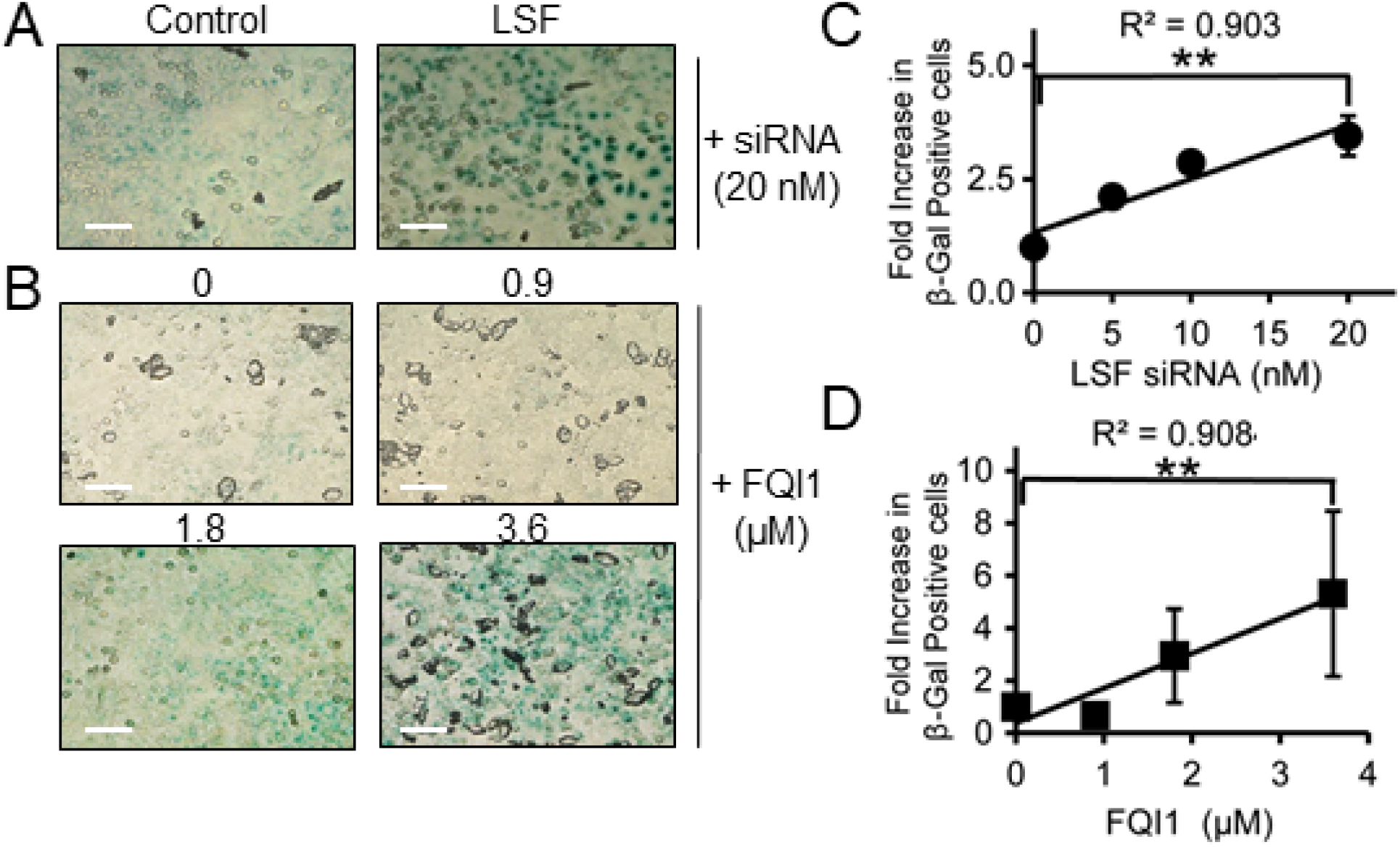
Inhibition of LSF activity induces cellular senescence. **A-B**. HeLa cells treated either with increasing concentrations of FQI1 (0=vehicle control) or with control or LSF siRNA were synchronized using a double thymidine block (protocols in Fig. 2A and 3A, respectively) and then fixed at 8 h after release from the second thymidine block and stained for β-galactosidase activity. Phase contrast images were taken at 20x magnification. Images shown are representative of three independent experiments. **C-D.** The correlation of increasing LSF siRNA concentrations (C) or increasing FQI1 concentrations (D) with the number β-galactosidase positive cells is depicted as a fold change compared to the control for each individual FQI1 and LSF siRNA concentration. The data reflect analysis of 75 cells per condition in each experiment, averaging over three independent experiments. Pearson correlation coefficients are indicated. Scale bars: 50 µm.

## Discussion

LSF is an oncogene in multiple cancer types, notably including hepatocellular carcinoma (1, 2, 20). Small molecule inhibitors directly targeting LSF inhibited hepatocellular carcinoma cell proliferation i*n vitro* and tumor growth *in vivo* with no signs of toxicity at doses required for tumor inhibition (15–17). Together, these data suggested that LSF is a promising therapeutic candidate for hepatocellular carcinoma patients, and likely for other cancer types. In order to confirm the key characteristic that anti-tumor effects of FQIs were due to specific targeting of LSF, we demonstrated here that a siRNA targeting LSF produced strikingly similar results to that of FQI1 treatment in all aspects, confirming specific targeting by the small molecule inhibitor. Furthermore, knockdown of the close LSF paralog, LBP1A, did not result in such mitotic defects. Thus, we conclude that LSF is the FQI1 target that is required for accurate and efficient mitotic progression in these cancer cells.

The primary consequence of LSF inhibition was the delay in progression through mitosis with condensed, but unaligned chromosomes. This can lead to multiple cellular outcomes. First, if unable to progress through normal metaphase alignment, mitotic slippage yields multinucleated cells, which lead to apoptosis, shown previously in hepatocellular carcinoma cell lines (15, 16), and here in HeLa cells as sub-G1 cellular DNA content (Fig. 1A, Supplementary Fig. S4). Second, less severe mitotic defects that permit cell division can also result in senescence after division, yielding cells with “2n” genomic DNA content unable to progress further through the cell cycle. Here, we demonstrate induction of senescence in a dose-dependent manner by treating with either LSF inhibitor or LSF-targeting siRNA during the synchronization protocol (Fig. 5).

The simplest interpretation for the mitotic defects observed upon LSF inhibition would be that, as a transcription factor, LSF directly regulates expression of mitotic regulators. Both FQI1 and LSF siRNA do result in downregulation of *AURKB* and *CDC20* expression, initially suggesting that the mitotic phenotypes could be caused by inhibition of Aurora kinase B and/or CDC20. Aurora kinase B inhibition leads to defects in kinetochore-microtubule attachment and cytokinesis, followed by multinucleation (39, 40), and knockdown of CDC20 results in an increase in mitotic time (28). However, upon deeper analysis, including demonstration of the onset of senescence in a subpopulation of the inhibited cells, it is unclear at this time as to whether lower AURKB and CDC20 levels cause, or rather are the consequence of, disruption of normal cell cycle progression. Nonetheless, their diminished expression provide molecular biomarkers for LSF inhibition *in vitro*. Importantly, our analysis conclusively demonstrated that alteration of cyclin B levels are unrelated to the FQI1-mediated mitotic phenotype, refuting a previously reported interpretation from experiments using asynchronous populations of hepatocellular carcinoma cells (16). We propose that the observed, elevated cyclin B protein levels in FQI1-treated versus control cells resulted simply from accumulation of cells in mitosis when treated with FQI1 for 12-24 hours, whereas the control cells continually cycled in the asynchronous populations. Furthermore, the CDK1/cyclin B inhibitors used in Rajasekaran et al. to attempt to rescue the mitotic defects are not specific to inhibiting CDK1 activity — cycloheximide influences many cell cycle processes, and Roscovitine also robustly inhibits the cell cycle regulator CDK2, thus complicating the interpretations made. Overall, these complications underscore the need to perform population-level experiments in synchronized cells, as shown here, when dissecting mitotic defects.

Since FQIs are specific for targeting LSF, and both these inhibitors and siRNAs targeting LSF can induce cell death or senescence in cancer cells *in vitro*, it is worthwhile to consider the targeting strategy for LSF inhibition in patients. Many cancer drug candidates target mitosis in an effort to exploit this key vulnerability of cancer cells. However, many such therapies have failed in trials, which may result from: (1) tumor escape, where pathway redundancy or evasive resistance in mammalian cells enables the tumor cell to escape the therapy (41, 42), or (2) low mitotic index where the drug half-life may not be long enough to suppress the target when cell division is triggered for any particular tumor cell (43, 44). As a target, LSF may have an advantage toward avoiding tumor escape. Inhibiting a transcription factor can target multiple pathways simultaneously, thus the likelihood that system redundancy would fully compensate is diminished. In addition, the issue of low mitotic index may be avoidable for the LSF inhibitors, since the apparent lack of toxicity in preclinical models may permit dosing in manners that generate sustained drug levels. Gene silencing based approaches may also provide a useful strategy to counter low mitotic index for hepatocellular carcinoma patients. The first RNAi drug, which uses a lipid nanoparticle to encapsulate and efficiently deliver siRNA to hepatocytes, was recently approved following robust and durable gene silencing over the 18-month pivotal study (45). Additionally, a ligand-based strategy to deliver LSF siRNA to hepatocytes may provide added benefit as recent human data using a triantennary *N*-acetylgalactosamine (GalNAc) mediated siRNA delivery system demonstrated robust knockdown of a hepatic target that was sustained for more than a year (46, 47). The target of GalNAc, asialoglycoprotein receptor (ASGR1) (22), is expressed in early stages and often in later stages of hepatocellular carcinoma (48), although whether tumors retain ubiquitous expression is not clear.

## Conclusions

The high specificity of FQI1 for LSF was confirmed by comparing cellular and molecular outcomes of small molecule inhibitors that eliminate LSF activity to those achieved following targeted LSF protein depletion using RNAi technology, Both mechanisms resulted in similar mitotic defects, followed by cellular death or senescence, proving that LSF regulates mitosis in cancer cells. Therefore, the anti-tumor activity of FQI1 in multiple preclinical models is explicitly due to loss of LSF activity. These findings support the candidacy of LSF targeting agents for treatment of hepatocellular carcinoma, as well as other cancers in which LSF is identified as an oncogene.

## Funding

The research was supported by: Alnylam Pharmaceuticals, Inc (JLSW, KF), Boston University (JLSW, UH, SES), BU Undergraduate Research Opportunity (MR), New England Biolabs, Inc. (HGC), and the NIH (R01 GM078240) to SES.

## Authors’ contributions

JLSW performed and interpreted all the experiments except the time lapse and ChIP experiments, and drafted the manuscript. KG performed the time lapse data acquisition and analysis. MPR generated the fluorescently tagged cells used for the time lapse study. HGC performed the ChIP experiment. PS generated RNA from mitotic shakeoff cells. HS and CSP analyzed the ChIP data. SES provided the LSF small molecule inhibitors. KF provided overall guidance and support. JS contributed to experimental design and interpretation oversight of the time lapse experiments. UH contributed to overall experimental design, interpretation of results, and revision of the manuscript. All authors approved of the final manuscript.

## Notes

#### Summary of Updates

The manuscript has been updated for the following reasons: 1) to highlight the advantages of using synchronized cell populations in studying the cellular consequences of LSF inhibition, 2) to now emphasize, with additional analysis in the Discussion, that a previous publication based on experiments in which asynchronously growing cells were treated with LSF inhibitors was incorrect in suggesting that elevation of cyclin B levels was the cause of the resulting mitotic arrest, 3) to reformat the previous bar graphs in order to include all data points, rather than averages, and 4) to revise the title, in order to more accurately reflect the major contributions of the study.

