## Supplementary Information&Figures for "Targeting the oncogene LSF with either the small molecule inhibitor FQI1 or siRNA causes mitotic delays with unaligned chromosomes, resulting in cell death or senescence"

#### Phase contrast and fluorescent microscopy

Cells were imaged using an Axiovert 40 CFL (Zeiss) microscope for both phase contrast and fluorescent imaging. Paraformaldehyde fixed cells were analyzed using a Zeiss Axioimager M1 microscope utilizing both 63x and 100x magnifications to analyze mitotic progression based on DNA staining by DAPI (Invitrogen) and by antibodies indicated below.

Immunofluorescence

Cells were fixed in paraformaldehyde, permeabilized with 0.1% Triton X-100, blocked with 1% Bovine Serum Albumin (Sigma Aldrich), and incubated with one or more primary antibodies containing 1% Bovine Serum Albumin overnight at 4°C. Primary antibodies included γH2AX (Cell signaling 9178S) at a 1:50 dilution, LSF (Abcam, ABE180), α-Tubulin (Abcam AB7750), or Aurora Kinase B (Abcam AB2254). Secondary antibodies included anti-rabbit Alexa 488 (Abcam AB150069) and anti-mouse Alexa 647 (Abcam AB150111).

Gene expression determination from mitotic cells

For isolation specifically of mitotic RNA, HeLa cells were synchronized to the G1/S border using a double thymidine block, as in Fig. 2A, adding FQI1 or vehicle at the indicated concentrations. At the second G1/S block, any loosely attached cells (e.g. mitotic or dying) were removed by a double shake-off procedure. The remaining cells were released in growth media containing 20 μM thymidine plus the respective FQI1 or vehicle concentrations. Seven hours after the release, mitotic shake-off was performed and repeated four times in 15 minute increments. After each shake-off, mitotic cells were immediately centrifuged and kept on ice. RNA from the combined mitotic cell pellets was purified using TrizolTM (Invitrogen), according to manufacturer's instructions.

#### Viability and cell counting

Cells were counted on the ViCell XR coulter counter from Beckman Coulter. The ViCell also assayed viability by measuring Trypan Blue permeability. Fifty individual fields per sample were assayed to determine total cell and total viable cell populations.

#### Chromatin immunoprecipitation analysis

A HEK293 cell line in which HA-tagged LSF is inducibly expressed was incubated in 5 μM RSL1 ligand for 24 hr, and harvested using the SimpleChIP Plus Enzymatic Chromatin IP Kit protocol (Cell Signaling Technology). Briefly, cells were fixed with formaldehyde.

Chromatin was digested with Micrococcal Nuclease into chromatin fragments containing 150-900 bp DNA, immunoprecipitated with IgG or antibody targeting HA (Abcam), and captured by Protein A magnetic beads. After reversing cross-links, 50 ng of purified DNA, mono- or di-nucleosome-sized, were used for library preparation using the NEBNext Ultra II DNA library Prep Kit for Illumina (NEB E7645S). Libraries were constructed and sequenced in separate lanes on a Illumina GAII platform to obtain 72 bp paired-end reads. Reads that mapped to the reference genome (hg19) were used to call statistically significant peaks using MACS (Model-based Analysis of ChIP-seq), modified to take into account the use of MNase-digested chromatin as starting material. Binding of LSF to peak regions was validated by qPCR, using the following primers. AURKB promoter: 5' GGGAGAGTAGCAGTGCCTTG 3'and CCAACGGACCCTCTGATCTA; AURKB upstream: 5' AATTAGCTGGATGTGGTGGC 3' and 5' CAAGCAATTCTTGTGCCTCA 3'.

### SUPPLEMENTARY FIGURES

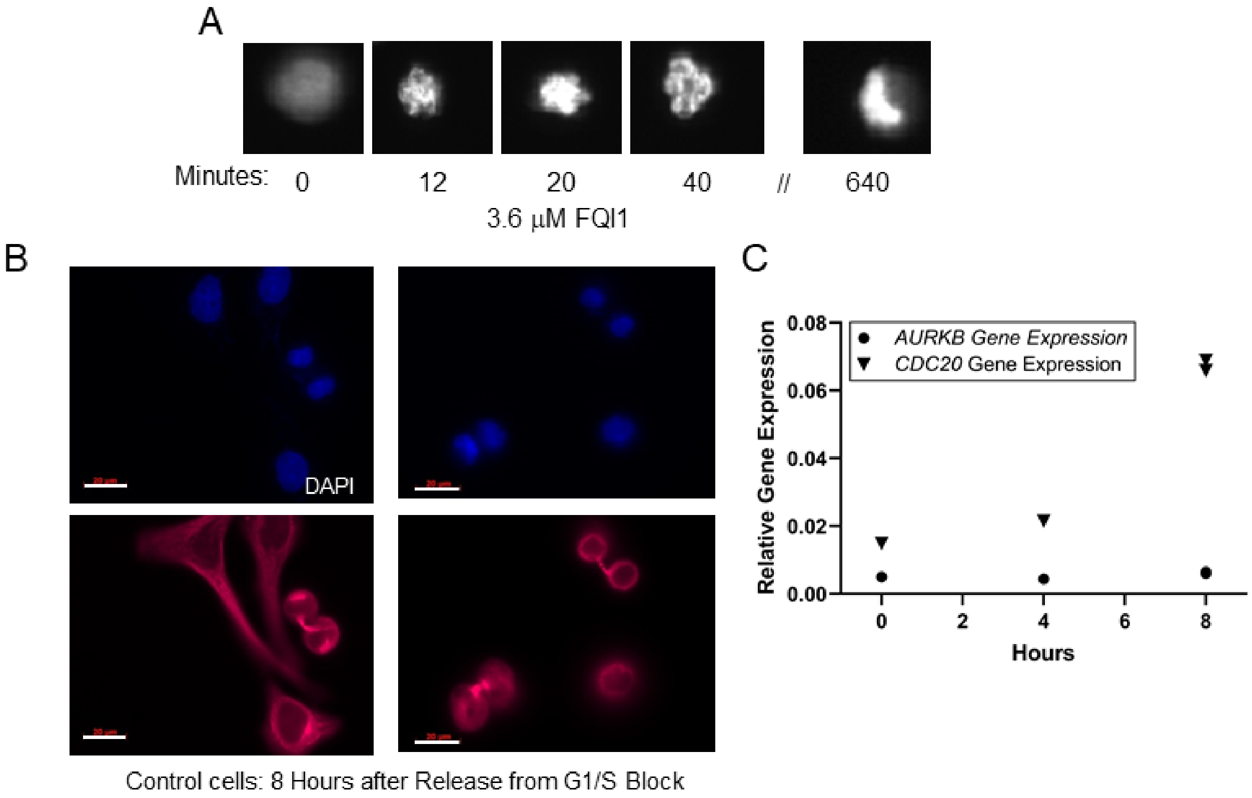

**Supplementary Figure S1. Characterization of synchronized HeLa cells and mitotic arrest with high FQI1 concentrations. A.** Representative images are shown of H2B-YFP-expressing HeLa cells treated with 3.6 µM FQI1. Images were taken during time lapse microscopy, with 0 minutes indicating the time frame at which the imaged cell underwent nuclear envelop breakdown. All cells that entered mitosis were arrested with condensed, but nonaligned chromosomes. **B.** Representative images from vehicle-treated HeLa cells synchronized using a double thymidine block (protocol in Fig. 2A) and fixed for staining at 8 hours after release from the G1/S block. Cells were stained for DNA using DAPI (Blue) and probed with an antibody detecting α-tubulin (Red). Scale bars: 20 µm. **C.** Relative gene expression of *AURKB* and *CDC20* in vehicle-treated, synchronized HeLa cells harvested at 0 (G1/S), 4 (S) and 8 (M) hours post release from a G1/S block. Both *AURKB* and *CDC20* levels were normalized to those of *GAPDH*. Data points derived from two independent experiments.

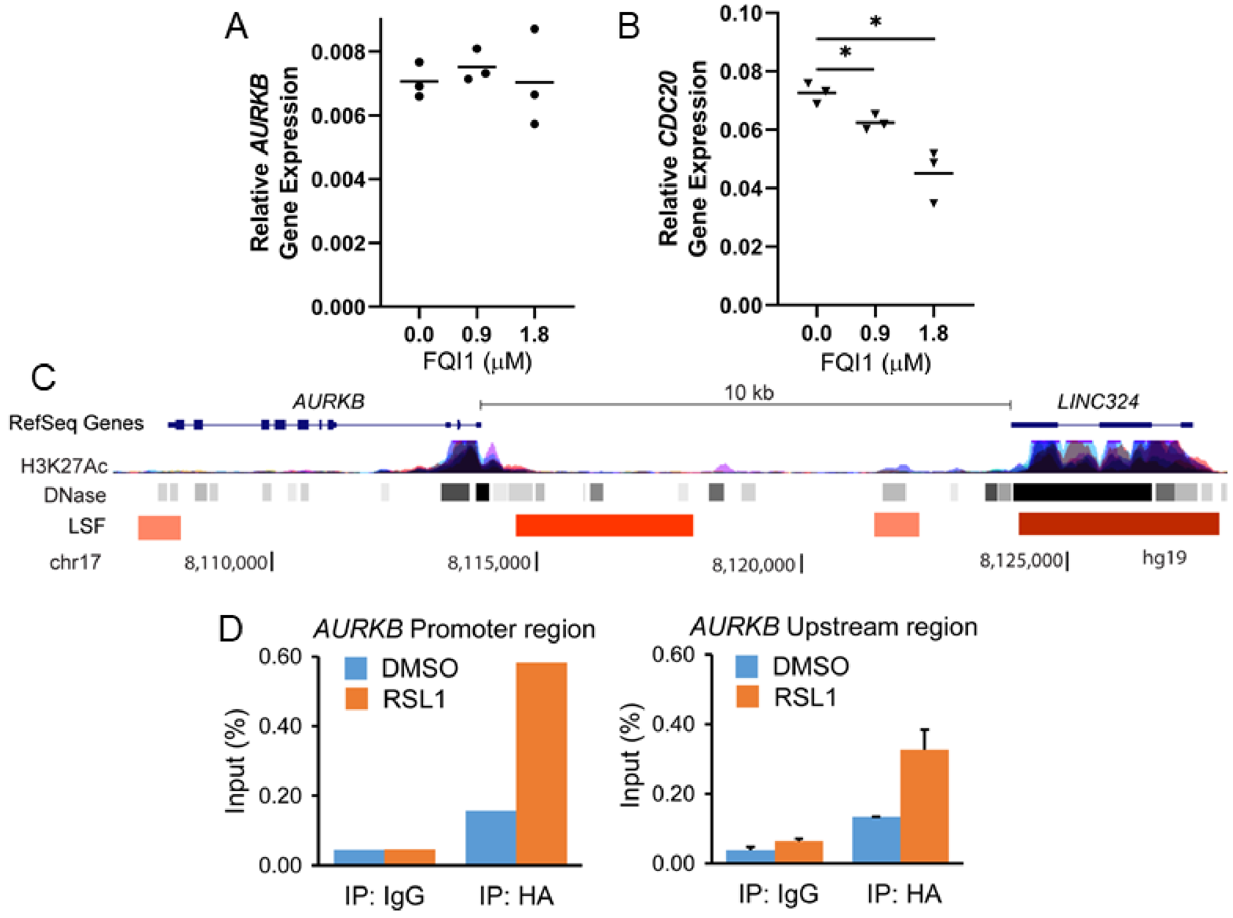

**Supplementary Figure S2. A.-B.** HeLa cells synchronized at the G1/S border with a double thymidine block were treated with or without FQI1 during the synchronization protocol, and were harvested in mitosis after release from the block by mitotic shakeoff. Multiple shakeoff populations were pooled, RNA isolated, and analyzed for levels of *AURKB* (**A**) and *CDC20* (**B**) transcripts, as normalized to those of *GAPDH*. Shown are data points and means from three independent experiments. *p=0.028 (**B**, 0.9 µM FQI1), p=0.038 (**B**, 1.8 µM FQI1). **C.** LSF ChIP-Seq data around the *AURKB* gene. Shown are genomic and composite chromatin features around *AURKB*, including H3K27 acetylation and DNase hypersensitive regions (http://genome.ucsc.edu/). Regions identified by ChIP-Seq as binding HA-LSF are indicated in red, with the more robust MACS peaks indicated in darker shades. **D.** LSF-binding sites surrounding the *AURKB* gene. Binding of induced HA-LSF was verified in independent biological samples by qPCR to the promoter region (left) and within the peak around -2800 upstream (right).

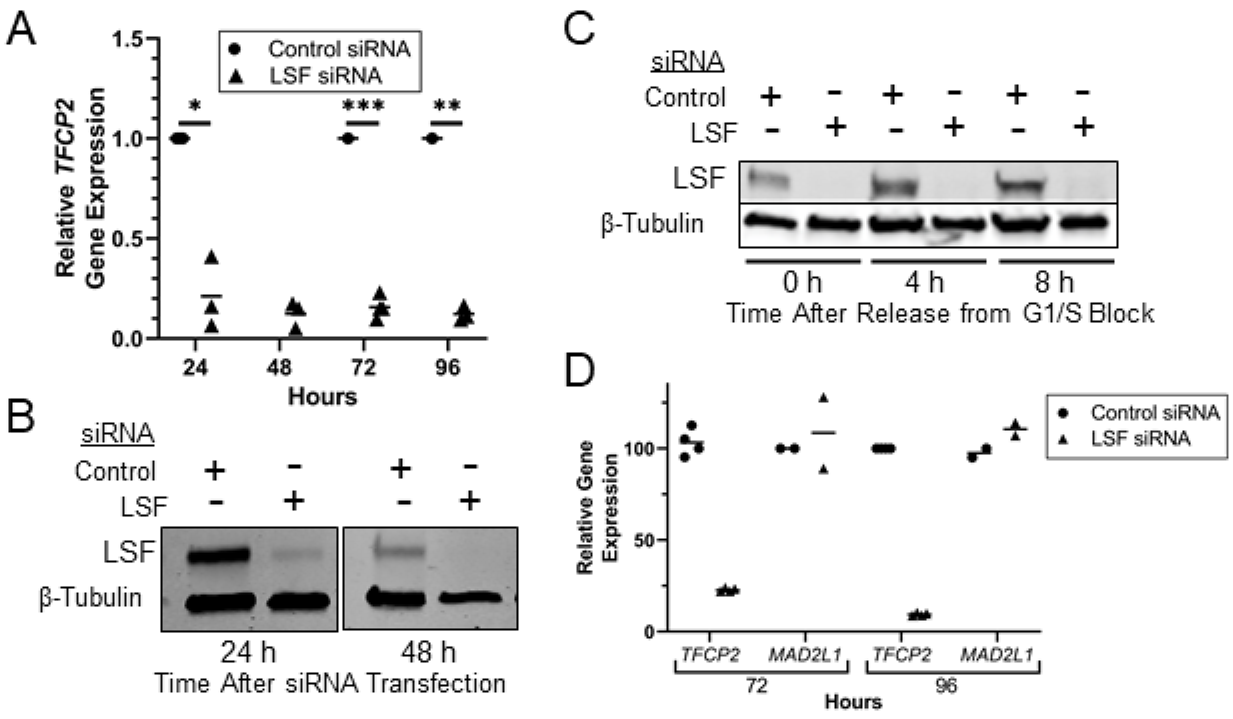

**Supplementary Figure S3. LSF RNAi-mediated knockdown is robust and durable.** siRNAs targeting LSF or a non-expressed target (control) were transfected into HeLa cells at 20 nM. **A.** Levels of *TFCP2* RNA (encoding LSF) from asynchronous cells at the indicated times after transfection were measured. Relative RNA levels, normalized to those of *GAPDH*, are then normalized to the levels in the control siRNA-treated cells. Shown are data points representing averages of technical duplicates from 3-4 independent experiments, with overall means indicated by horizontal lines. *p=0.017; **p=0.0023 (unpaired T test); ***p=0.0008 (unpaired T test). **B-C.** Representative immunoblots of LSF and β-tubulin are shown from lysates harvested at various times during the cell synchronization and release protocol (see Fig. 3A). **B.** Lysates were collected at 24 and 48 hours post transfection during the double thymidine block procedure. **C.** Lysates were collected following release from the final thymidine block for 0, 4 and 8 hours. **D.** Asynchronous HeLa cells were transfected with control or LSF siRNA, and harvested at the indicated times. Relative *TFCP2* (encoding LSF) and *MAD2L1* RNA levels are shown, normalized to those of *GAPDH*. Data points and means derived from 2-4 independent experiments, each with 2-4 technical replicates.

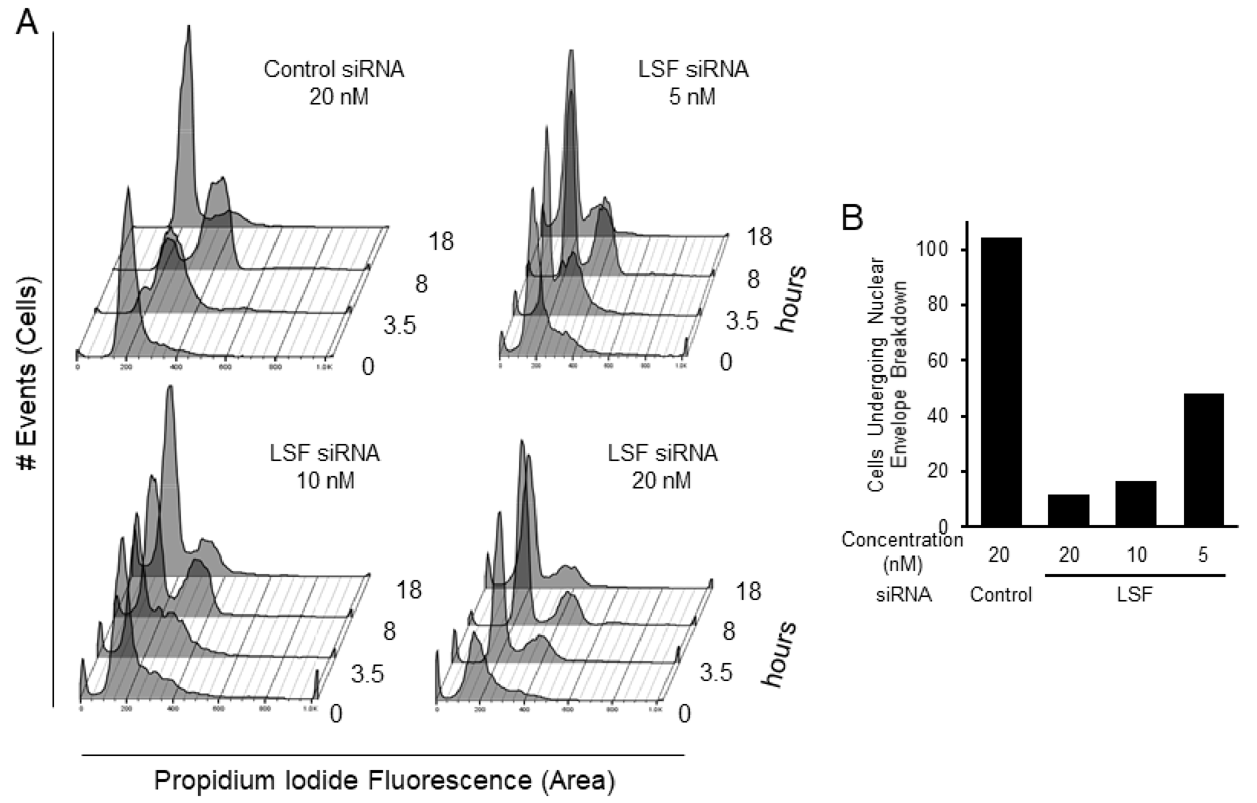

**Supplementary Figure S4. Synchronization of HeLa cells after LSF knockdown reveals two distinct phenotypes: a static 2n DNA population and a population progressing from 2n to 4n DNA content transitioning to a subG1 population. A.** HeLa cells transfected with control or LSF siRNA at the indicated concentrations were synchronized using the double thymidine block protocol (see Fig. 3A). At the indicated times after release from the final block, cells were fixed and stained with propidium iodide for analysis of cellular DNA content by flow cytometry. Data are representative of more than 4 experiments. **B.** Dose-dependent decrease in cells entering mitosis with decreasing levels of LSF activity in synchronized HeLa cells. H2B-YFP-expressing HeLa cells were transfected with LSF siRNA at the indicated concentrations or with a control siRNA. Cells were synchronized using a double thymidine block and then released (Fig. 3A). Upon release the cells were analyzed by time-lapse microscopy. Shown are the number of imaged cells that entered mitosis in each condition, based on nuclear envelope breakdown (NEB) within the 10-12 hour time frame of the experiment. A minimum of 100 cells were counted in the control cells.

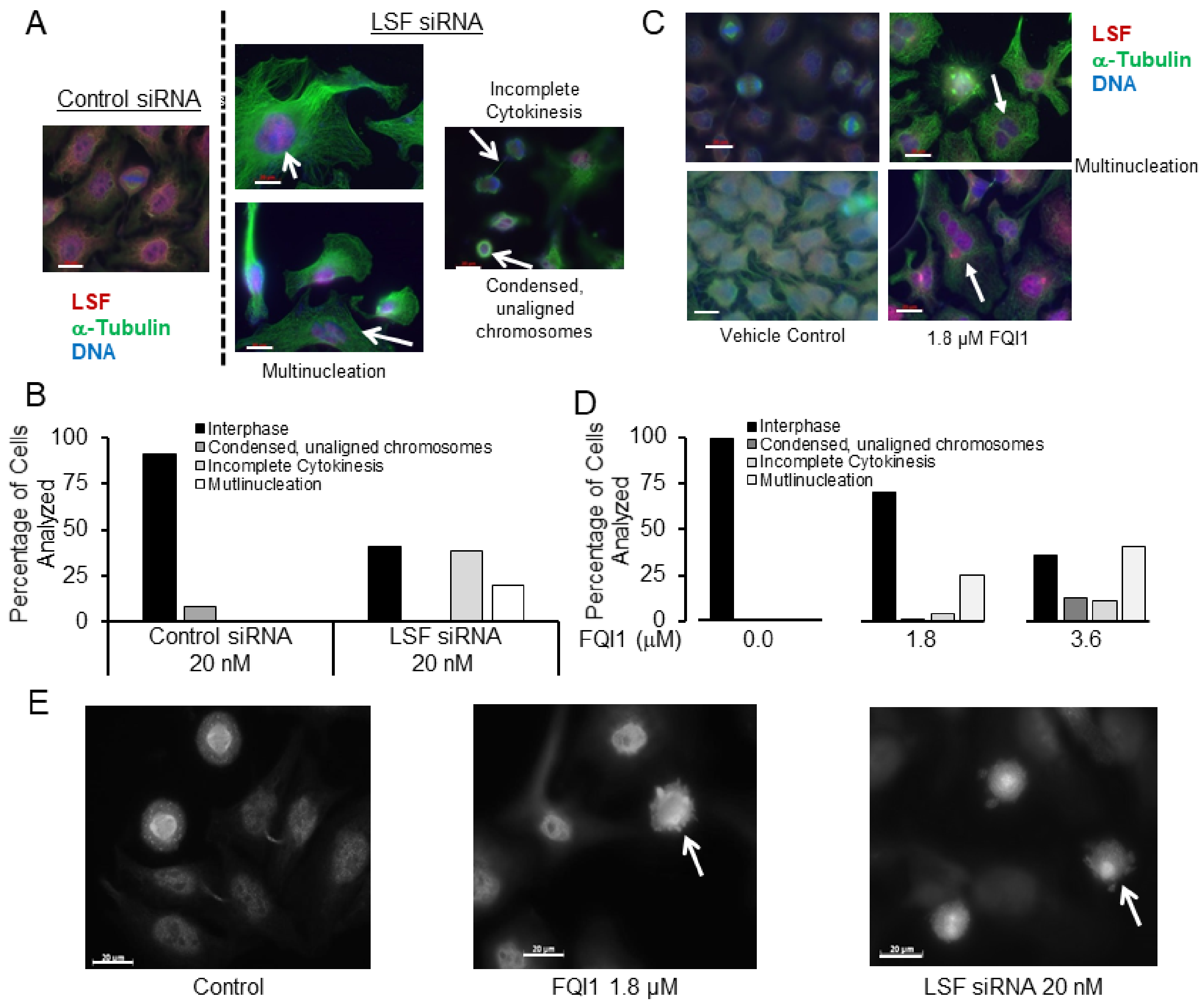

**Supplementary Figure S5. LSF inhibition by FQI1 or siRNA results in multi-nucleation and lack of complete cytokinesis. A.** Following transfection with LSF or control siRNA at 20 nM, HeLa cells were synchronized using a double thymidine block, and then released to progress through the cell cycle. Cells were fixed and probed with an anti-LSF (red) and anti-α-tubulin (green) antibody and stained with DAPI (blue). Images were analyzed at 16 hours post release from the block, when control cells progressed back into G1 phase, at 63x magnification. Representative images are shown. Arrows point to cells with the phenotypes indicated next to the images. Scale bars: 20 μm. **B.** Quantitation of 89-102 individual cells within each group from the experiment in panel **A** were analyzed from both the control and LSF siRNA groups for the indicated, observed cellular phenotypes. **C.** HeLa cells were synchronized using a double thymidine block at the G1/S border following incubation with vehicle control or FQI1 at concentrations of 1.8 μM and then released to continue with cell cycle progression. Cells were fixed and probed with anti-LSF (red) and anti-α-tubulin (green) antibodies, as well as being stained for DNA with DAPI (blue). Images were analyzed at 16 hours post release at 63x magnification. Approximately 100 individual cells were analyzed within each group. Representative images are shown. Arrows point to multinucleated cells. Scale bars: 20 μm. **D.** Quantitation of 98-107 individual cells within each group from the experiment in panel **C** were analyzed for the indicated, observed cellular phenotypes. **E.** Synchronized HeLa cells were treated with either 20 nM of LSF siRNA or 1.8 μM of FQI1. Cells were collected 8 hours after release from the final G1/S block and fixed for immunofluorescent analysis with an anti-α-tubulin antibody. Arrows indicate the cytoskeletal protrusions. Scale bars: 20 μm.

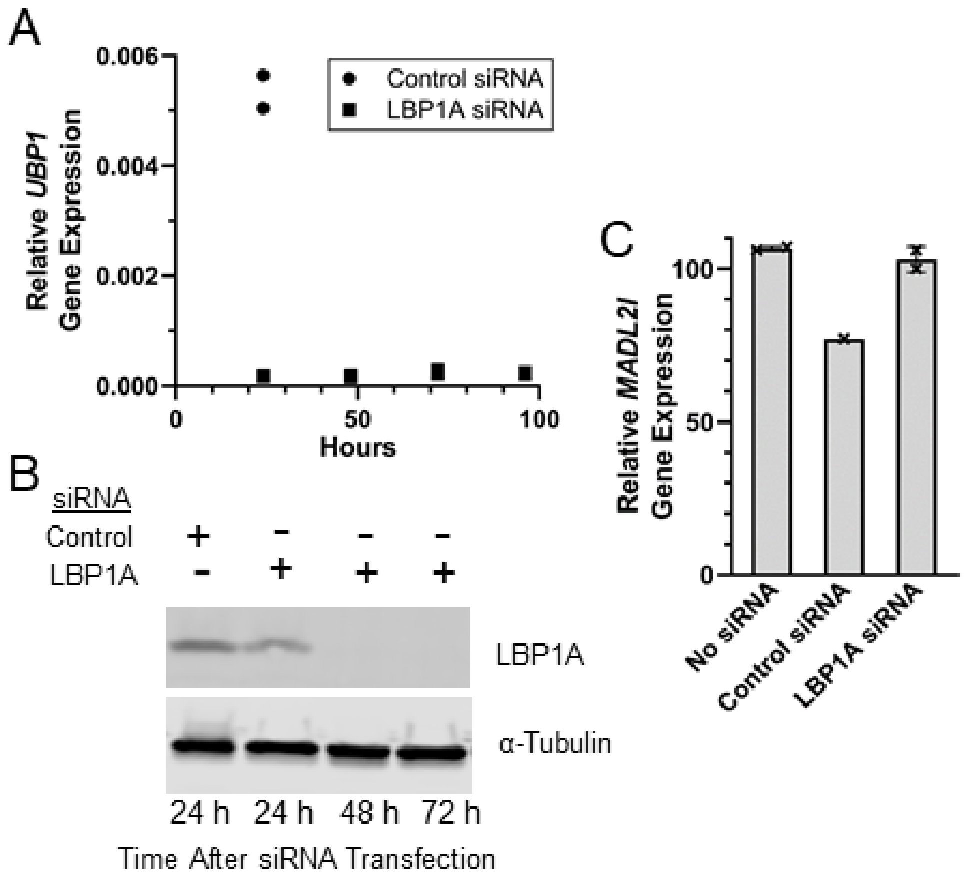

**Supplementary Figure S6. LBP1A RNAi-mediated knockdown is robust and durable.** siRNAs targeting LBP1Aa or a non-expressed target (control) were transfected into HeLa cells at 20 nM. **A.** Levels of *UBP1* RNA (encoding LBP1A) from asynchronous cells were measured at the indicated times after transfection. Relative RNA levels, normalized to those of *GAPDH*, are then normalized to the levels in the control siRNA-treated cells. Data points are from pairs of technical replicates. **B.** Representative immunoblots of LBP1A and α-tubulin are shown for lysates harvested at the indicated times during the cell synchronization protocol using a double thymidine block (see Fig. 3A). **C.** Asynchronous HeLa cells were either not transfected (None), or transfected with control siRNA or LBP1A siRNA and harvested 72 hours later. Relative *MAD2L1* RNA levels are shown, normalized to those of *GAPDH. MAD2L1* expression is depicted as a percentage of levels in control cells treated with no siRNA (just the vehicle, phosphate-buffered saline). Data points with means and SD are shown from 2 independent experiments.

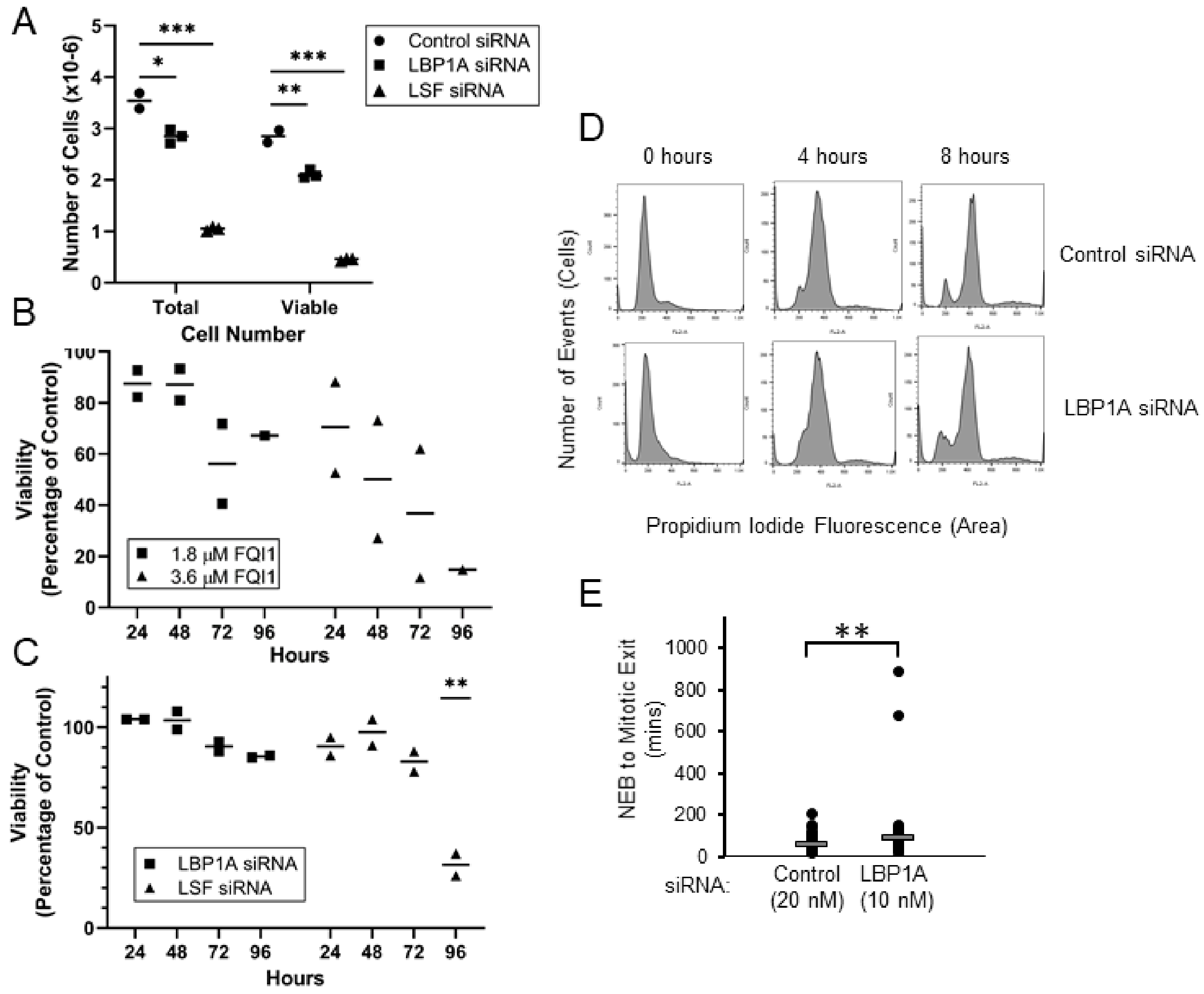

**Supplementary Figure S7. LBP1A knockdown in HeLa cells results in reduced cell number, but minimal impact on viability or cell cycle progression. A.** Total cell number was measured simultaneously to viability measurements on asynchronous HeLa cells treated with 20 nM of control siRNA or siRNAs targeting either LSF or LBP1A at 120 hours post transfection. Data points are technical replicates and means from a single experiment; results are representative of three independent experiments. **B.** Asynchronous HeLa cells were treated with increasing concentrations of FQI1 and analyzed at the indicated times for cell viability by the Promega MTT assay, performed as the manufacturer's instructions. The data are represented as the percentage of viability compared to that in the control. Data represent averages with standard error of the mean calculated from three independent experiments. **C.** Asynchronous HeLa cells were transfected with 20 nM of control siRNA or siRNAs targeting either LSF or LBP1A. Cells were analyzed at the indicated times for viability by the Promega MTT assay. The data are represented as the percentage of viability, compared to that of the control siRNA. Data points and means are derived from 2 independent experiments. **p=0.0064 (LSF vs. control siRNA, unpaired T test). **D.** HeLa cells were synchronized following transfection with siRNAs targeting LBP1A or a non-expressed control using a double thymidine block. Cells were analyzed for cellular DNA content by flow cytometry at the indicated time points after release from the G1/S block to continue cell cycle progression. **E.** H2BYFP-labeled HeLa cells were synchronized with a single thymidine block following transfection with a siRNA targeting either LBP1A or a non-expressed gene as a control. Cells were imaged using time lapse microscopy. Mitotic time was determined by measuring time from nuclear envelope breakdown (NEB) to anaphase. Mitotic times (mean time in minutes +/- standard error, n) for 20 nM control siRNA and 10 nM LBP1A siRNA were: 57.9 +/- 2.8, 101; and 86.6 +/- 10.6, 100; respectively. ** p=0.009.
